# Lis1 relieves cytoplasmic dynein-1 auto-inhibition by acting as a molecular wedge

**DOI:** 10.1101/2022.10.10.511666

**Authors:** Eva P. Karasmanis, Janice M. Reimer, Agnieszka A. Kendrick, Jennifer A. Rodriguez, Joey B. Truong, Indrajit Lahiri, Samara L. Reck-Peterson, Andres E. Leschziner

**Author notes:** Co-corresponding authors. S.L.R.P; A.E.L. Co-first authors (Eva P. Karasmanis, Janice M. Reimer, Agnieszka A. Kendrick). Equal contribution (Samara L. Reck-Peterson, Andres E. Leschziner).

## Abstract

Cytoplasmic dynein-1 transports many intracellular cargos towards microtubule minus ends. Dynein is autoinhibited and undergoes conformational changes to form an active complex, consisting of one or two dynein dimers, the dynactin complex and activating adaptor(s)^1,2^. The Lissencephaly 1 gene, *LIS1*, is genetically linked to the dynein pathway from fungi to mammals and is mutated in patients with the neurodevelopmental disease lissencephaly^3–5^. Lis1 is required for active dynein complexes to form^6–10^, but how it does so is unclear. Here, we present a structure of two dynein motor domains with two Lis1 dimers wedged in-between. The contact sites between dynein and Lis1 in this structure, termed “Chi”, are required for Lis1’s regulation of dynein in *Saccharomyces cerevisiae* in vivo and the formation of active human dynein–dynactin– activating adaptor complexes in vitro. We propose that this structure represents an intermediate in dynein’s activation pathway, revealing how Lis1 relieves dynein’s autoinhibited state.

---

Cytoplasmic dynein-1 (dynein) is the main minus-end directed microtubule motor responsible for transporting vesicles, protein complexes, RNAs, organelles, and viruses along microtubules. Dynein also positions nuclei and the mitotic spindle during mitosis and meiosis^2^. Dynein is conserved across eukaryotes, with the exception of flowering plants and some algae, and requires numerous binding partners and regulators to function^2,11^. Mutations in dynein or its regulators cause neurodevelopmental and neurodegenerative diseases in humans, while homozygous deletion of dynein is embryonically lethal in mice^12,13^. In contrast, in the yeast *S. cerevisiae*, dynein and its regulators are conserved but non-essential, providing an important model system to study dynein’s mechanism and function.

Dynein is a 1.4 MDa complex consisting of a dimer of two motor-containing heavy chains composed of a ring of six AAA+ (ATPase associated with various cellular activities) domains, two intermediate chains, two light intermediate chains, and two copies of three different light chains^14^. The dynein AAA+ ring is dynamic and ATP binding and hydrolysis in AAA1 regulates dynein’s binding to and movement on microtubules. Opening and closing of dynein’s ring is coupled to movements of dynein’s mechanical element, the “linker”, and rearrangements in dynein’s stalk and buttress are responsible for controlling the affinity of the microtubule-binding domain for microtubules^15^ (Fig. 1a).

**Figure 1.**
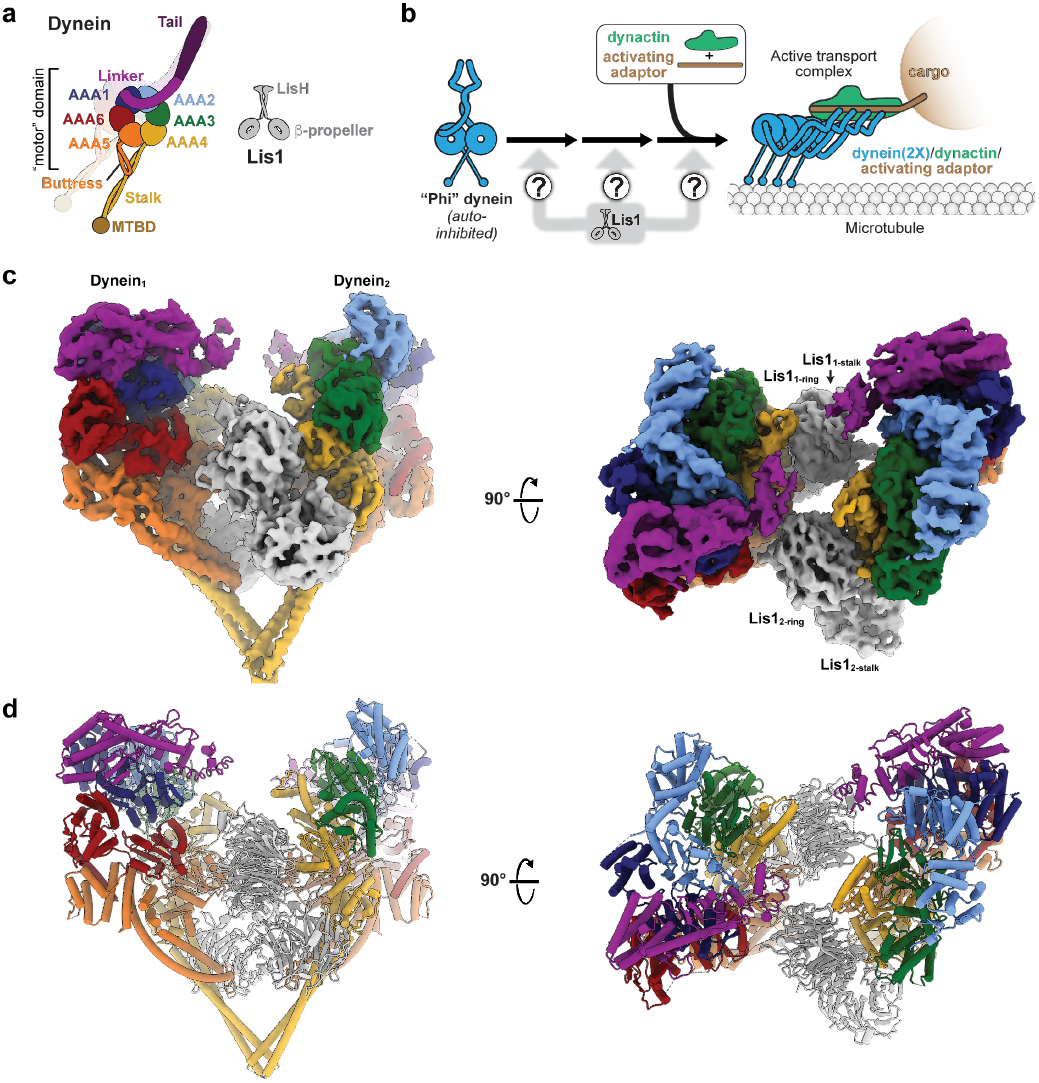
Structure of the Chi dynein-Lis1 complex. **a**. Cartoons of dynein’s and Lis1’s domain organization. Major elements mentioned in the text are labelled. **b**. Schematic of dynein activation. Activation of dynein requires the relief of its autoinhibited conformation (Phi), and the formation of an active complex containing the dynactin complex and an activating adaptor. While Lis1 is known to be involved in this process, how it is involved is unknown. **c**. Cryo-EM map of the Chi dynein-Lis1 complex, shown in two orientations. The different Lis1 β-propellers that bind to previously identified sites on dynein are labeled. **d**. Model of the Chi dynein-Lis1 complex, shown in the same two orientations as the map in (c).

Dynein is proposed to exist largely in an autoinhibited ‘Phi’ conformation in cells, which has been observed in vitro^16–18^ (Fig. 1b). In contrast to Phi dynein, which only contains the dynein subunits, active dynein complexes can be > 4 MDa and consist of one or two dynein dimers, the 1.1 MDa dynactin complex, and an activating adaptor that mediates the interaction between dynein and dynactin and links dynein to its cargos^19,20^ (Fig. 1b). Dynein must undergo major conformational changes to transition from the autoinhibited Phi form to these active complexes^1,18,21^. Some activated dynein complexes contain two dynein dimers, which move faster than complexes containing a single dynein dimer (Fig. 1b)^21–24^, and can also include a second activating adaptor^1^. Many putative activating adaptors have been described and about a dozen of these have been shown to activate dynein motility in vitro, including members of the Hook, Bicaudal D (BicD), and Ninein families^25–27^. Yeast dynein requires dynactin and the presumed activating adaptor Num1 for function in vivo as well^28–30^.

Dynein function in vivo also requires Lis1, which is an essential positive regulator of dynein, as shown by genetic studies in many organisms^31–36^. Like dynein, Lis1 is conserved from yeast to mammals. Lis1 is a dimer, with each monomer consisting of an N-terminal small dimerization domain, an alpha helix, and a C-terminal β-propeller following a flexible linker^37,38^ (Fig. 1a). Lis1 is the only dynein regulator known to bind directly to dynein’s AAA+ motor domain, with two known binding sites: one on dynein’s motor domain between AAA3 and AAA4 (site_ring_) and the other on dynein’s stalk, a long anti-parallel coiled coil that leads to dynein’s microtubule binding domain (site_stalk_)^39–42^. In humans, mutations in Lis1 (*PAFAH1B1*) or the dynein heavy chain (*DYNC1H1*) cause the neurodevelopmental disease lissencephaly and other malformations of cortical development^4,5,13^. Recent studies show that Lis1 has a role in forming active dynein complexes^22,24,40,43^. Human dynein-dynactin-activating adaptor complexes form more readily in vitro in the presence of Lis1 and move faster on microtubules due to the recruitment of a second dynein dimer to the complex^22,24^. In *S. cerevisiae* and the filamentous fungus *Aspergillus nidulans*, mutations in dynein that block the formation of the autoinhibited Phi conformation can partially rescue Lis1 deletion or mutations, suggesting that Lis1 may activate dynein by relieving autoinhibition^34,43,44^. On the other hand, the inability of Phi blocking mutations to fully rescue Lis1 loss of function phenotypes suggests that Lis1 may have additional roles in regulating dynein beyond relieving the autoinhibited Phi conformation^34,40,43^.

Despite the data summarized above, how Lis1 relieves dynein autoinhibition remains unknown. Previously, we determined a high-resolution cryo-EM structure of the *S. cerevisiae* dynein motor domain (carrying a E2488Q mutation in the Walker B motif of its AAA3 domain) in the presence of ATP-Vanadate and bound to two yeast Lis1 β-propellers, most likely coming from the same Lis1 dimer^40^. Although this dataset appeared to consist primarily of particles containing one dynein motor domain bound to one Lis1 dimer, we revisited it to look for other species. To our surprise, additional data processing revealed several 2D class averages that contained two dynein motor domains, even though the dynein we used for cryo-EM sample preparation was engineered to be monomeric. Using this subset of particles, we solved a structure of two dynein motor domains complexed with two Lis1 dimers to 4.1 Å. We termed this structure ‘Chi’ because this new dynein conformation resembles the Greek letter Chi, and because Chi follows Phi in the Greek alphabet

(Fig. 1c and 1d; Supplementary Video 1; Extended Data Fig. 1; Table 1). In our structure, two dynein motor domains, each bound by a Lis1 dimer, come together through novel dynein-Lis1 interactions and the closed conformation of dynein’s ring and the characteristic bulge in the stalk near its contact site with the buttress indicate a weak microtubule binding state^45^. The overall structure of each dynein-Lis1 complex is very similar to the structures we solved of individual dynein motor domains bound to a Lis1 dimer, also in a weak microtubule binding state^39,40^.

**Table 1:** Cryo-EM data information and model validation. CryoEM data collection parameters, reconstruction information and model refinement statistics.

|  | Dynein<sup>E2488Q</sup> bound to Lis1 in Chi conformation | Dynein<sup>E2488Q</sup> bound to Lis1 in Chi conformation, symmetry expansion |
| --- | --- | --- |
| **Models** |  |  |
| PDB | 8DZZ | 8E00 |
| EMDB | EMD-27810 | EMD-27811 |
| **Data collection and processing** |  |  |
| Microscope | Titan Krios | Titan Krios |
| Camera | Gatan K2 Summit | Gatan K2 Summit |
| Voltage (kV) | 300 | 300 |
| Electron exposure (e-/Å<sup>2</sup>) | 58.3 | 58.3 |
| Defocus range (μm) | 2 – 2.7 | 2 – 2.7 |
| Pixel size (Å) | 1.31 | 1.31 |
| **Reconstruction** |  |  |
| Symmetry imposed | C2 | C1 |
| Initial particle images (no.) | 561 397 | 561 397 |
| Final particle images (no.) | 23 385 | 46770 |
| Micrographs collected (no.) | 2378 | 2378 |
| Map resolution (Å) (0.143 FSC threshold) | 4.1 | 3.6 |
| **Model Refinement** |  |  |
| Initial model used (PDB code) | 8E00 | 7MGM |
| Map-to-model resolution (0.5 FSC threshold) (Å) | 4.4 | 3.9 |
| Map sharpening *B* factor (Å<sup>2</sup>) | 106.5 | 90.5 |
| Model to Map Fit |  |  |
| CC mask | 0.80 | 0.82 |
| CC volume | 0.79 | 0.81 |
| CC peaks | 0.76 | 0.67 |
| *Model composition* |  |  |
| Non-hydrogen atoms | 49 638 | 24 808 |
| Protein residues | 6112 | 3055 |
| Ligands | 8 | 4 |
| *B* factors (Å<sup>2</sup>) |  |  |
| Protein | 154.9 | 94.1 |
| Ligand | 132.4 | 77.6 |
| *R.m.s. deviations* |  |  |
| Bond lengths (Å) | 0.005 | 0.005 |
| Bond angles (°) | 1.080 | 1.102 |
| *Validation* |  |  |
| MolProbity score | 1.10 | 0.94 |
| Clashscore | 3.11 | 1.84 |
| Poor rotamers (%) | 0.18 | 0.25 |
| *Ramachandran* |  |  |
| Favored (%) | 98.46 | 98.41 |
| Allowed (%) | 1.51 | 1.59 |
| Disallowed (%) | 0.03 | 0 |

Dynein’s autoinhibited Phi conformation is characterized by interactions between its two motor domains that make them point in opposite directions (they are effectively “cross-legged”) (Fig. 1b and 2a), which prevents the motor from being able to bind to microtubules^18^. The Phi conformation is incompatible with the binding of Lis1 to dynein, as Lis1 bound to one dynein clashes with the second dynein in Phi (Extended Data Fig. 2a).

**Figure 2.**
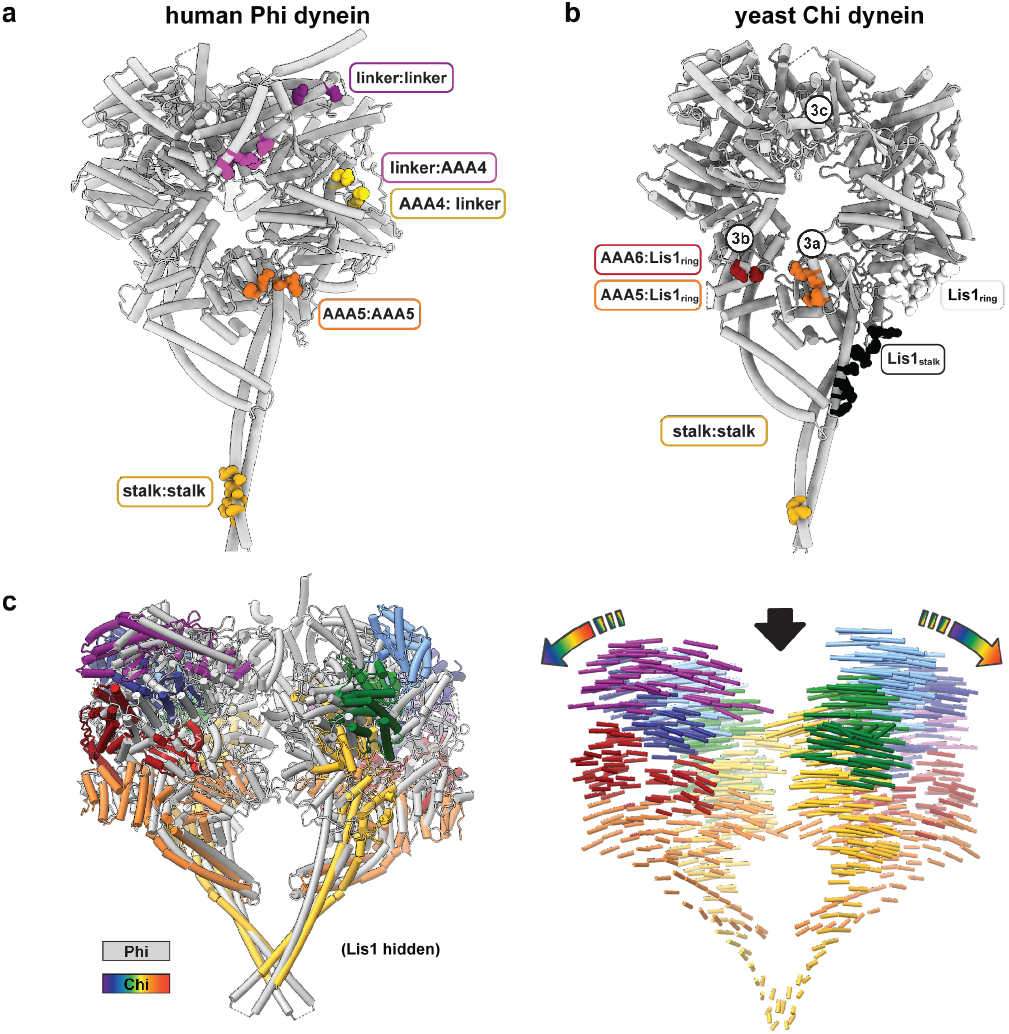
Comparison of dynein in Chi and Phi. **a**. Interfaces in human dynein involved in stabilizing Phi. **b**. Interfaces in yeast dynein involved in stabilizing Chi. Note that all but one (stalk:stalk) of the interfaces in Chi also involve Lis1, which we do not show in this panel. The circled labels (3a, 3b, 3c) refer to panels in Figure 3 where interfaces are shown in detail. **c**. Overlay of dynein in Phi (grey) and Chi (rainbow). As in panel (b), we omitted Lis1 in the Chi structure for clarity. The left panel shows the two structures superimposed. The right panel shows interatomic vectors linking equivalent alpha carbons in Phi and Chi. The length of the cylinders is proportional to the displacement of that atom between the two structures.

In Chi, the Lis1s bound to the dynein motor domains act as wedges that keep the two dyneins apart, preventing most of the interactions that stabilize Phi (Fig. 2a and 2b, Extended Data Figure 2b). The one exception is a contact between dynein’s stalk helices, which acts as a hinge where Phi opens to accommodate Lis1 and transition to Chi (Fig. 2a and 2b). The dynein-dynein Phi contacts that are disrupted in Chi are replaced by new contacts between Lis1 bound to site_ring_ on one dynein and the AAA5 (Fig. 3a) and AAA6 (Fig. 3b) domains of the opposite dynein. These Chispecific interactions involve residues on Lis1’s β-propeller that are different from those involved in all previously characterized Lis1-dynein and Lis1-Lis1 contacts (Fig. 3d-g; Supplementary Video 1). The Lis1 bound to site_stalk_ does not interact with the opposite dynein molecule in Chi (Fig. 1c and 1d).

**Figure 3.**
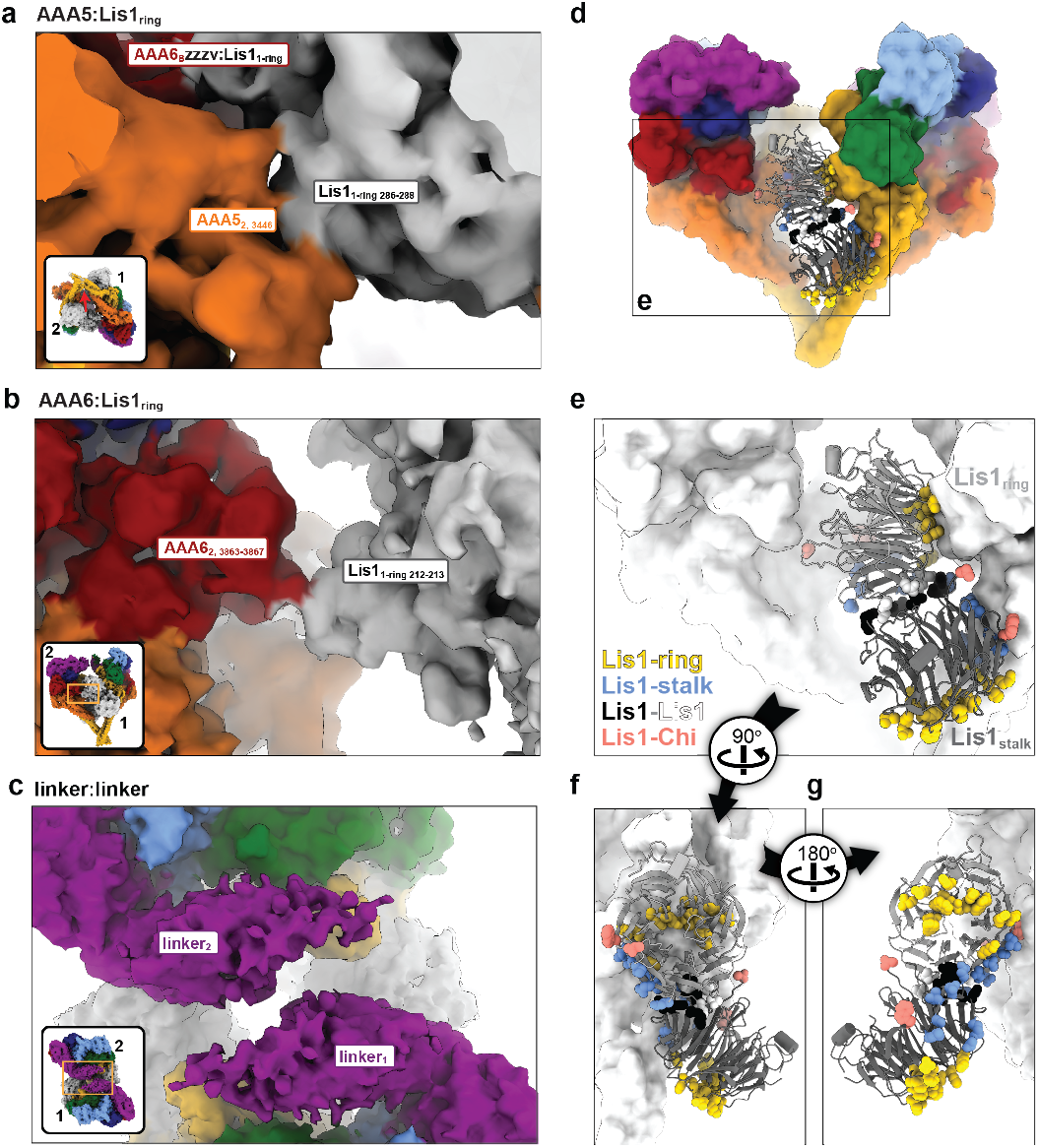
Comparison of dynein in Chi and Phi. **a-c**. Chi is stabilized by three novel interfaces: AAA5:Lis1_ring_ (a); AAA6:Lis1_ring_ (b), and a linker:linker interaction (c). Figure 2b shows the location of these interactions in the context of full dynein. All three panels show the cryo-EM map colored by domain, with Lis1 in grey. The insets show the region of Chi that is highlighted in the main panel. **d-g**. Location of interaction interfaces in the Lis1_ring_-Lis1_stalk_ dimer. **d**. Our model of Chi with dynein shown in surface representation (filtered to 8Å) and Lis1 in ribbon diagram. The area within the square is enlarged in (e). **e**. Residues involved in the four different Lis1 interactions (binding of Lis1 to site-ring and site-stalk, the Lis1-Lis1 dimer interface, and the interactions stabilizing Chi) are shown in sphere representation and colored by interface. Dynein is shown in white. **f**. The Lis1 dimer is viewed facing the dynein monomer with which the Lis1-ring and Lis1-stalk interactions are formed. The dynein in front was removed for clarity. **g**. The Lis1 dimer is viewed from the other side, facing the dynein monomer with which the Chi-stabilizing interactions are made. The dynein in front was removed for clarity.

The differences between Phi and Chi include changes in the conformation of the linker domain (Extended Data Fig. 3). In Phi dynein, the linker is bent and docked on AAA2/AAA3 (Extended Data Fig. 3a), which is the same conformation seen in dynein alone in its “pre-power stroke” state15. In contrast, in Chi, the linker is disengaged from AAA2/AAA3 and shifted towards the opposing dynein molecule (Fig. 3c and Extended Data Fig. 3b and 3c); this accommodates the increased distance between the motor domains brought about by the presence of Lis1 (Fig. 2c). This change in the linker conformation in turn causes a shift in the interface mediating the linker-linker interaction (Fig. 3c and Extended Data Fig. 3d-f); however, the resolution of the linker domain is too low to determine the exact nature of the residues involved.

The major features of the Chi structure—the disruption of most Phi-stabilizing interactions, a shift in the linker-linker interface, and the dependence of these on the presence of Lis1 and new interactions between Lis1 and dynein—suggest that Chi is an early intermediate in the activation of dynein that provides a mechanistic explanation for the role of Lis1 in this process. We set out to test this hypothesis and validate our structural model in vivo using *S. cerevisiae* and in vitro using recombinant human dyneindynactin-activating adaptor complexes.

We first sought to determine if the new contact sites between Lis1 and dynein at AAA5 (Fig. 4a) and AAA6 (Fig. 4b) that form Chi are important for the dynein pathway in vivo. For this, we turned to experiments in S. cerevisiae. In yeast, dynein functions to align the mitotic spindle, such that upon the completion of mitosis both the mother and daughter cell inherit a nucleus^28,30,46–48^. Deletion of dynein and dynactin subunits, as well as Lis1 causes nuclear segregation defects that are readily quantifiable as binucleate cells^31^. To disrupt the Chi-specific interactions (Fig. 4a and 4b) we mutated Lis1’s Asn 213 to Ala (Lis1N213A) and Trp 288 to Asp (Lis1W288D), or both (Lis1N213A, W288D) in the endogenous locus of yeast Lis1 (*PAC1*). Western blots of strains containing FLAG-tagged versions of the mutants confirmed that Lis1N213A, Lis1W288D and Lis1N213A, W288D were expressed at wild type levels (Extended Data Fig. 4a and 4b). We found that all three mutants exhibited an increase in the percentage of cells containing two nuclei, similar to cells where Lis1 was deleted (Fig. 4c). This suggests that the dynein-Lis1 interactions involved in forming Chi are required for Lis1’s regulation of dynein in vivo.

**Figure 4.**
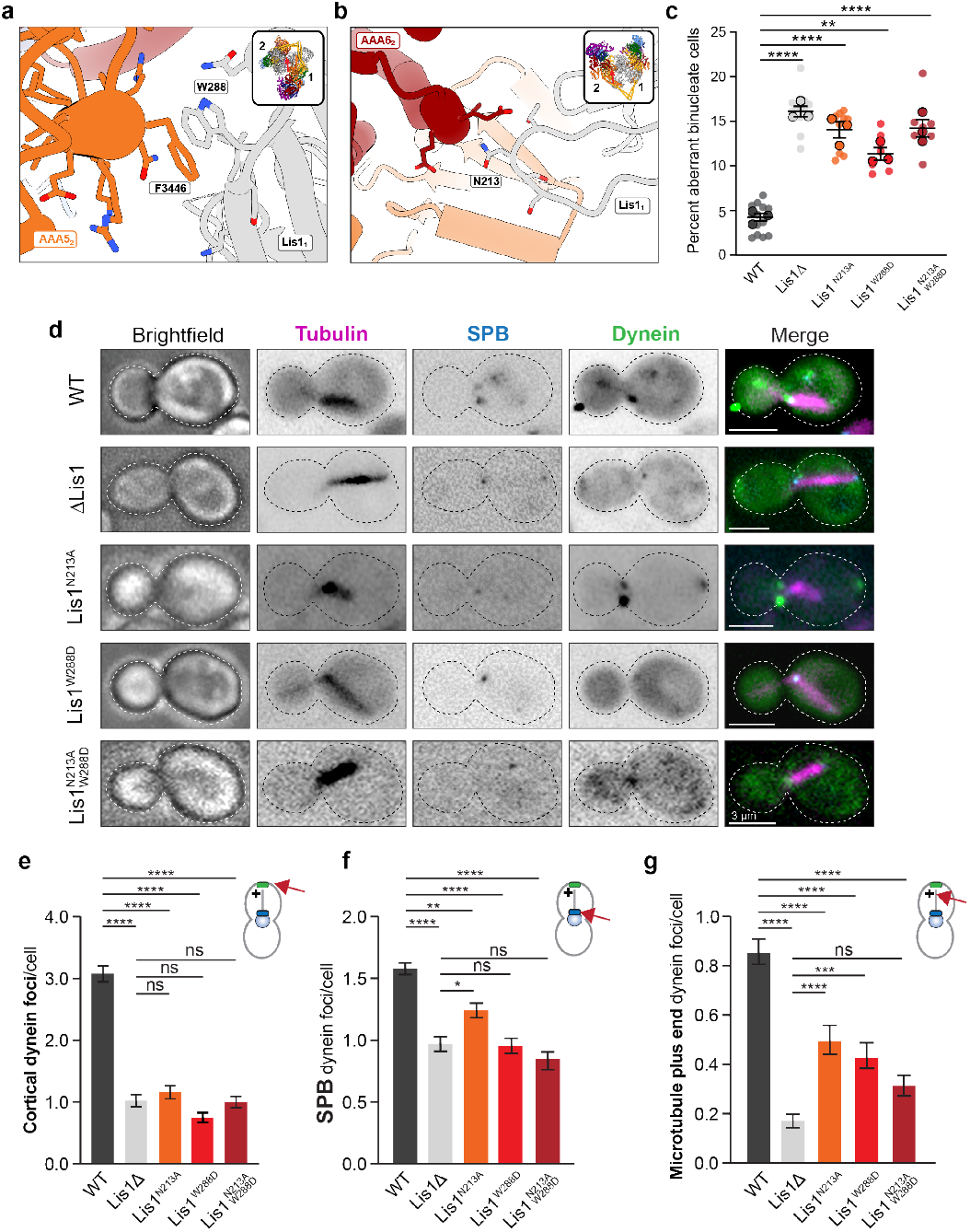
Role of Chi in dynein’s function in *S. cerevisiae*. **a**. The yeast Chi AAA5-Lis1_ring_ interface. Trp 288 in Lis1 was mutated to Asp. **b**. The yeast Chi AAA6-Lis1_ring_ interface. Asn 213 was mutated to Ala. **c**. Quantitation (mean ± s.e.m.) of the percentage of cells displaying an aberrant binucleate phenotype for wild type (WT, dark gray), Lis1 deletion (light gray), Lis1^N213A^ (orange), Lis1^W288D^ (red) and Lis1^N213A, W288D^ (maroon). n = 3 independent experiments, with 3-4 replicates per experiment, with at least 200 cells per condition. Statistics were generated using a One-way ANOVA with Tukey’s multiple comparisons of each mean test. **** WT and ΔLis1 p<0.0001, WT and Lis1^N213A^ p<0.0001, ** WT and Lis1^W288D^ p=0.0001, **** WT and Lis1^N213A, W288D^ p<0.0001. Differences not noted are not significant. **d**. Example images of endogenous dynein localization (DYN1-3xGFP) in dividing yeast cells expressing a fluorescently tagged SPB (*SPC110-tdTomato*) component and tubulin (*TUB1-CFP*). Scale bars are 3 μm. **e-g**. Quantification of dynein localization. Bar graphs show the average number of dynein foci per cell localized to (e) the cortex, (f) SPB, and (g) microtubule plus ends in WT, Lis1Δ, Lis1^N213A^, Lis1^W288D^ and Lis1^N213A, W288D^ yeast strains. Statistical analysis was performed using a one-way ANOVA with Tukey’s multiple comparison of means test; ****, p<0.0001; ***, p=0.0002; **, p=0.0018; *, p=0.025; ns (not significant), p=0.1825>0.9999. n = 120 cells per condition.

Next, we wanted to determine how the Lis1-dynein interactions involved in forming Chi contribute to dynein’s localization in yeast cells (Fig. 4d-g). Yeast dynein assembles into fully active complexes with dynactin and Num1, the presumed yeast dynein activating adaptor, at the cell cortex^28,49^. From this site, active dynein pulls on spindle pole body (SPB)-attached microtubules to align the mitotic spindle, which ultimately favors equal segregation of nuclei between mother and daughter cells. Dynein reaches the cortex by first localizing to microtubule plus ends, either via kinesin-dependent transport or through cytosolic recruitment^50–52^. Dynein also localizes to the SPB, where microtubule minus ends are found51. Lis1 is required for dynein’s localization to all three sites^30,31,53^. To assess how the mutants that target the new interactions between dynein and Lis1 in Chi affect dynein localization, we introduced Lis1N213A, Lis1W288D and Lis1N213A, W288D at the endogenous PAC1 locus. In these strains, dynein, alpha-tubulin and a spindle pole body component were also tagged with a fluorescent protein (DYN1-3xGFP, TUB1-CFP and SPC110-tdTomato, respectively) (Fig. 4d)^31,39^ and used to measure the number of dynein foci that were found at the cell cortex (Fig. 4e), SPBs (Fig. 4f), or microtubule plus ends (Fig. 4g) in each cell. All three mutants showed a striking reduction in the number of dynein foci at the cell cortex, similar to a Lis1 deletion strain (Fig. 4e). Because the cortex is the site of dynein-dynactin-Num1 assembly, these data suggest that the Chi conformation is important for the assembly of active dynein complexes in vivo. All three mutants also significantly affected the ability of dynein to localize to microtubule plus ends. One mechanism dynein uses to reach microtubule plus ends is via kinesin transport in a complex that also contains Lis1 and Bik1^30,54,55^. Thus, it is possible that this transport complex uses the novel dynein-Lis1 interactions we identified in Chi, or potentially the Chi conformation itself. Together, our data show that the contacts we observe between dynein and Lis1 in the Chi structure are required for active dynein complex assembly and function in vivo.

To assess the importance of the Chi conformation in the assembly of active human dynein-dynactin-activating adaptor complexes at the molecular level, we next turned to reconstituting the complexes using human proteins. In vitro, human Lis1 enhances the formation of dynein-dynactin-activating adaptor complexes containing two dynein dimers, which move faster than the single-dynein dimer complexes that form in the absence of Lis1^22,24^. To do this, we first had to identify Chi-disrupting mutations in human Lis1 predicted to be equivalent to the ones we identified in yeast. Directly overlaying our recent structure of a human dynein-Lis1 complex^56^ onto each motor domain of yeast Chi did not work, as Lis1 bound to site_ring_ is too far away from the opposite dynein to form the Chi-stabilizing interactions with AAA5 and AAA6 (Fig. 5a). This difference is due to human Lis1 having shorter peripheral loops and being slightly rotated relative to yeast Lis1^56^. To circumvent this problem, we modeled human Chi by bringing the motor domains closer together while maintaining the stalk-stalk interaction that functions as the hinge. Using this model, we made two sets of human Lis1 mutations: Lis1N203A, D205A, Y225A to disrupt both the Lis1-AAA5 and Lis1-AAA6 interfaces, and Lis1N203A, D205A, D245A to disrupt the Lis1AAA6 interface alone. Two of the sites we mutated in human Lis1, N203A and D245A, are also conserved in yeast, while Y225 corresponds to the W288 we mutated in the yeast Lis1 (Extended Data Fig. 5a).

**Figure 5.**
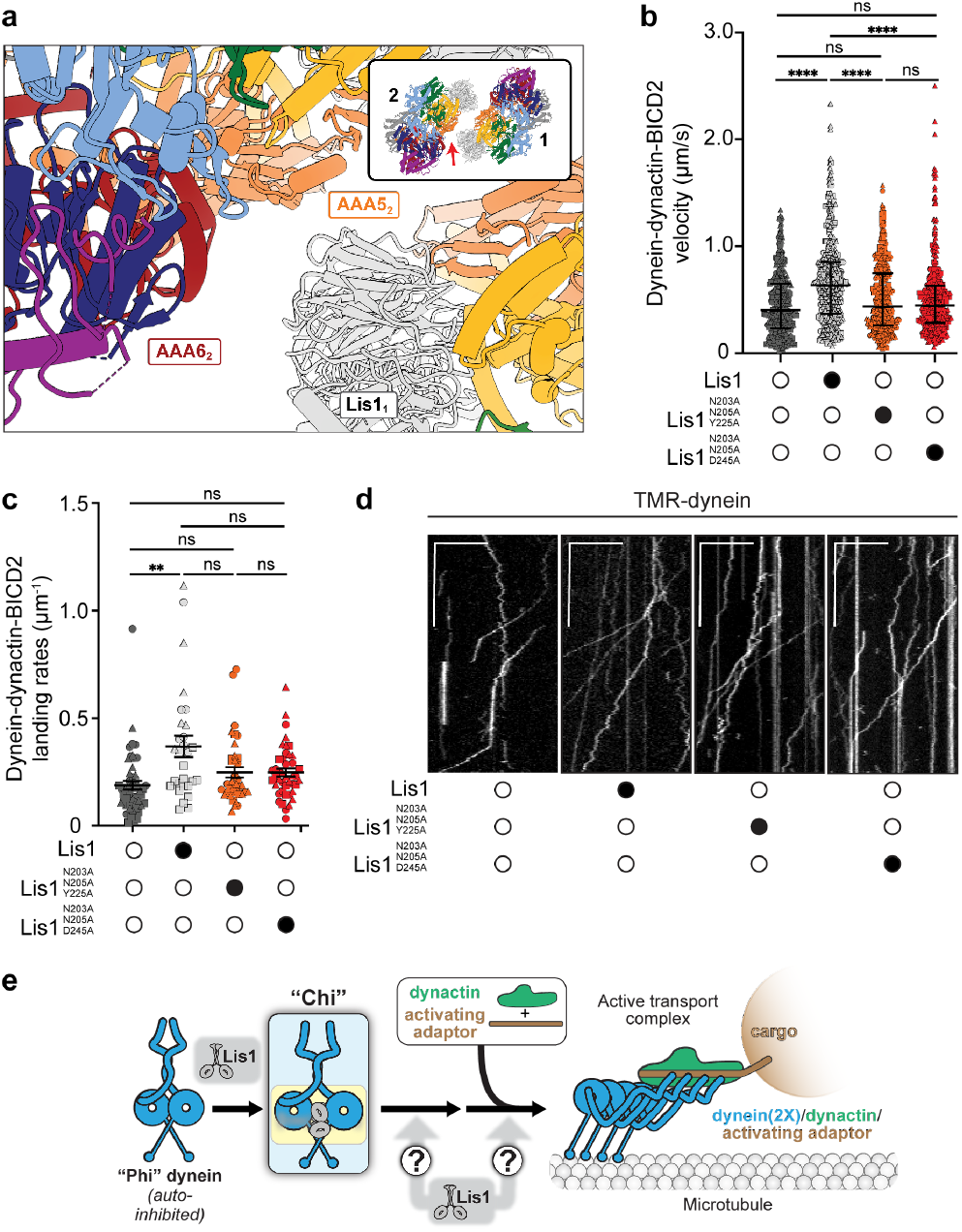
Role of Chi in human dynein’s ability to move on microtubules. **a**. A model of the human Chi AAA6_2_ and AAA5_2_-LIS1_1_ interface. The arrow in the inset indicates the area highlighted in the main panel. **b**. Single-molecule velocity of TMR-dynein-dynactin-BICD2 complexes in the absence (white circles) or presence (black circles) of different human Lis1 constructs. The data points are represented as triangles, circles and squares, corresponding to single measurements within each technical replicate (no Lis1, n = 367; Lis1, n = 408; Lis1^N203A, D205A, Y225A^, n = 360; Lis1^N203A, D205A, D245A^, n = 384). ns, not significant, **** p<0.0001, One-Way Anova with Tukey-post test. **c**. Landing rates (median ± interquartile range) of TMR-dynein-dynactin-BICD2 complexes in the absence (white circle) or presence (black circle) of different unlabeled human Lis1 constructs. The data points are represented as triangles, circles, and squares, corresponding to single measurements within each technical replicate. (no Lis1, n = 52; Lis1, n = 29; Lis1^N203A, D205A, Y225A^, n = 38; Lis1^N203A, D205A, D245A^, n = 43). ns, not significant, p=0.9272, One-Way Anova with Brown-Forsythe test. **d**. Representative kymographs from single-molecule motility assays with purified TMR-dynein-dynactin-BICD2 in the absence (white circle) or presence (black circle) of different human Lis1 constructs. Scale bars, 10 μm (*x*) and 40 s (*y*). **e**. The schematic of Chi hypothesis for how Lis1 relieves dynein autoinhibition.

Next, we purified these mutant human Lis1 constructs (Extended Data Fig. 5b) and examined their ability to activate human dynein-dynactin complexes containing the BICD2 activating adaptor in single-molecule motility assays. In these assays, activation of motility is read out by observing an increase in dynein complex velocity in the presence of Lis1, which results from the enhanced formation of dynein complexes containing two dynein dimers^21,22,24^. As we showed previously, pre-incubation of dynein-dynactinBICD2 with 300 nM wild type human Lis1 increased the velocity and landing rates of these complexes (Fig. 5b-d)^24^. In contrast, there was no significant difference between dynein velocity and landing rates when dynein-dynactinBICD2 complexes were pre-incubated with the two human Lis1 mutants (Lis1N203A, D205A, Y225A and Lis1N203A, D205A, D245A) as compared to complexes that were formed in the absence of Lis1 (Fig. 5b-d). We also observed a modest increase in the number of diffusive events in the presence of Lis1N203A, D205A, D245A (Extended Data Fig. 5c). These data indicate that the novel dynein-Lis1 contact sites found in the Chi structure are important for human Lis1’s role in forming the activated human dynein–dynactin–activating adaptor complex.

Based on the data presented here, we propose that Chi is an intermediate state in the dynein activation pathway, providing the first structural and mechanistic explanation for how Lis1 relieves dynein autoinhibition. We propose that the wedging of Lis1 between the two dynein motor domains primes dynein for binding dynactin and activating adaptor protein(s).

The Chi conformation has important implications for dynein activation. In dynein’s autoinhibited Phi state, the “tails”, which precede the motor domain, make many contacts with each other until they reach their “neck” region, where they move away from each other before rejoining again at the linker domain. Switching to the active dynactinbound form requires the tail to undergo a large conformational change that turns the loose two-fold symmetry present in Phi into the translational (“parallel”) symmetry dynein adopts when bound to dynein and an activating adaptor (Fig. 5e). This rearrangement involves breaking the linker-linker contacts, as well as many of the interactions between the tails and associated chains. We hypothesize that the Chi conformation primes dynein for binding dynactin and an activating adaptor protein by stabilizing an intermediate conformation of the tail. In Chi, binding of Lis1 forces the linkers apart, which would cause some of these interactions to break. This would pull the dynein heavy chains apart from each other, disrupting the neck and potentially transmitting conformational changes all the way up the tail.

## Acknowledgements

We thank the Nikon Imaging Center at UC San Diego and the Cryo-EM Facility at UC San Diego. We thank Zaw Htet for early work preparing and collecting data on the Lis1 sample used for CryoEM here. We also thank our funding sources: EPK is funded by a Jane Coffin Childs Postdoctoral Fellowship; JMR is a Merck Fellow of the Damon Runyon Cancer Research Foundation, DRG-2370-19; AAK is supported by American Cancer Society PF18-190-01-CCG. AL’s lab by NIH R01 GM107214 and R35 GM145296; and SRP’s lab by the Howard Hughes Medical Institute and NIH R35 GM141825. This paper was typeset with the bioRxiv word template by @Chrelli: www.github.com/chrelli/bioRxiv-word-template.

## Competing interest statement

The authors declare that they have no competing interests.

### Materials and Methods

#### Electron microscopy sample preparation

Grids were prepared and imaged as previously described^40^. Briefly, monomeric yeast dyneinE2488Q was randomly biotinylated using water-soluble Sulfo ChromaLink biotin and then dialyzed into TEV buffer. Grid samples containing 150 nM dyneinE2488Q, 650 nM Lis1, 1.2 mM ATP, and 1.2 mM Na3VO4 were applied to streptavidin affinity grids2–4 and vitrified^57,58^.

#### Electron microscopy image collection and processing

Details for image collection and initial processing are reported here^40^. Following particle picking in crYOLO5^59^, particles were extracted and binned to 3.93 Å/pixel in Relion 3.0^60^. Multiple rounds of 2D classification were carried out first in Relion 3.0 and subsequently in cryoSPARC^61^ to remove bad particles. Some class averages showed particles with more than one dynein molecule. Using particles from all good 2D classes, initial ab-initio reconstructions failed to reconstruct a volume containing more than one dynein molecule. Particles belonging to 2D class averages that potentially contained more than one dynein molecule were manually selected and used in an ab-initio reconstruction that resulted in an initial map for chi dynein bound to Lis1. This volume was used together with our previously published map of monomeric dyneinE2488Q bound to Lis1 (EMDB 23829) to separate monomeric dyneinE2488Q from chi dynein using heterogeneous refinement. We performed one additional round of heterogeneous refinement to further remove any monomeric dynein molecules. Using our final particles, we ran non-uniform refinement with C2 symmetry and optimized per-group CTF params enabled to give us a 4.1 Å map.

#### Model building and refinement

Symmetry expansion followed by local refinement was used to improve the overall resolution of the motor domain to 3.6 Å. The yeast dyneinE2448Q(Lis1)2 model (PDB 7MGM) was docked, and rigid body fit into the map using Phenix real space refine^62^. Discrepancies between the model and map were fixed manually in COOT6^63^ and then refined using a combination of Phenix real space refine7 and Rosetta Relax (v.13)^64^. Once the monomer model was finished, it was placed in the original map and refined using a similar strategy.

#### Cloning, plasmid construction, and mutagenesis

The pDyn1 plasmid (pACEBac1 expression vector containing insect cell codon-optimized dynein heavy chain [DYNC1H1] fused to a His-ZZ-TEV tag on the amino-terminus and a carboxy-terminal SNAPf tag (New England Biolabs) and the pDyn2 plasmid (the pIDC expression vector with codon optimized DYNC1I2, DYNC1LI2, DYNLT1, DYNLL1, and DYNLRB1) were a gift from Andrew Carter (LMB-MRC, Cambridge, UK). The pDyn1and pDyn2 plasmids were recombined in vitro with a Cre recombinase (New England Biolabs) to generate the pDyn3 plasmid. The presence of all six dynein chains was verified by PCR. The pFastBac plasmid with codon-optimized human full-length Lis1 (PAFAH1B1) fused to an amino-terminal HisZZ-TEV tag was a gift from Andrew Carter (LMB-MRC, Cambridge, UK). The BICD2s construct (amino acids 25-398) fused to sfGFP on the amino-terminus and inserted into a pET28a expression vector was obtained as described previously^26^. All mutations were made via site-directed mutagenesis (Agilent).

#### Yeast Strains

The *S. cerevisiae* strains used in this study are listed in Supplementary Table 1. The endogenous genomic copy of PAC1 (encoding for Lis1) was deleted using PCR-based methods as previously described^65^. In brief, K. lactis URA3 with homology arms complementary to regions upstream and downstream of the PAC1 genomic locus was generated using PCR. This fragment was transformed into a strain with the preferred genetic background using the lithium acetate method^66^ and screened by colony PCR. Point mutants were generated using QuikChange site-directed mutagenesis (Agilent) and verified by DNA sequencing. Mutated fragments were re-inserted into the klURA3 strains to reintroduce the mutated PAC1 gene. Positive clones (klURA3-) were selected in the presence of 5-Fluorootic acid, screened by colony PCR, and verified by DNA sequencing.

#### Nuclear segregation assay

Single colonies were picked and grown at 30°C. Log-phase *S. cerevisiae* cells growing at 30°C were transferred to 16°C for 16 hr. Cells were fixed with 75% ethanol for 1 h, sonicated for 5 sec at 40% amplitude and mounted in media containing DAPI. Imaging was performed using a ×100 Apo TIRF NA 1.49 objective (Nikon, Plano Apo) on a Nikon Ti2 microscope with a Yokogawa-X1 spinning disk confocal system, MLC400B laser engine (Agilent), Prime 95B back-thinned sCMOS camera (Teledyne Photometrics), and a piezo Z-stage (Mad City Labs). Imaging was blinded. The percentage of aberrant binucleate cells was calculated as the number of binucleate cells divided by the sum of wild-type and binucleate cells. This assay was repeated three completely independent times spanning over a year, with three to four biological replicates each time, with at least 200 cells counted for each replicate per condition.

#### Live-cell imaging

Single colonies were picked and grown at 30°C. Log-phase live *S. cerevisiae* cells were mounted on a thin agarose pad made from SC media pressed between two glass slides. Live cells genetically modified to express fluorescently labeled DYN1-3XGFP, CFP-TUB1, and SPC110-tdTomato were imaged using a Yokogawa W1 confocal scanhead mounted to a Nikon Ti2 microscope with an Apo TIRF 100 × 1.49 NA objective (Nikon, Plano Apo). The microscope was run with NIS Elements using the 488 nm 515 nm and 561 nm lines of a six-line (405 nm, 445 nm, 488 nm, 515 nm, 561 nm, and 640 nm) LUN-F-XL laser engine and Prime95B cameras (Photometrics). The number of DYN1-3XGFP foci localizing to the spindle pole body (SPB), microtubule plus end, and cell cortex was outlined as regions of interest in Fiji^67^, recorded and analyzed for three replicates of at least 120 cells for each sample.

#### *S. cerevisiae* immunoprecipitations and western blots

Log-phase *S. cerevisiae* cells grown at 30°C were pelleted at 4000 × g, resuspended in water and flash frozen in liquid nitrogen. Liquid nitrogen-frozen yeast cell pellets were lysed by grinding in a chilled coffee grinder, resuspended in Dynein-lysis buffer (30 mM HEPES [pH 7.4], 50 mM potassium acetate, 2 mM magnesium acetate, 1 mM EGTA, 10% glycerol, 1 mM DTT) supplemented with 1 mM Pefabloc, 0.2% Triton, cOmplete EDTA-free protease inhibitor cocktail tablet (Roche), and 1 mM Pepstatin A (Cayman Chemical Company) and spun at 50,000 × g for 1 hr. The protein concentration of the clarified supernatants was quantified using a Bradford Protein Assay (Bio-Rad), and equal amounts of clarified lysates were incubated with anti-FLAG M2 Affinity Gel (Sigma) overnight at 4°C. Beads were washed with Dynein-lysis buffer, boiled in SDS sample buffer, and loaded onto a NuPAGE Bis-Tris gel (Invitrogen). Gels were transferred to a PVDF membrane that was blocked with PBS-T (PBS1X and 0.1% Tween-20) containing 5% milk and 1% BSA for 1 hr at room temperature and blotted with a rabbit anti-FLAG antibody (1:3000; Proteintech 20543-1-AP) overnight at 4°C. Membranes were then incubated with a goat-anti-rabbit IRDye 6800RD secondary antibody (LI-COR) and were scanned in a ChemiDoc Imaging system (Bio-Rad).

#### Protein expression and purification

##### S. cerevisiae dynein

Protein purification steps were done at 4°C unless otherwise indicated. S. *cerevisiae* dynein constructs were purified from *S. cerevisiae* using a ZZ tag as previously described^68^. Briefly, liquid nitrogen-frozen yeast cell pellets were lysed by grinding in a chilled coffee grinder and resuspended in Dynein-lysis buffer supplemented with 0.1 mM Mg-ATP, 0.5 mM Pefabloc, 0.05% Triton, and cOmplete EDTA-free protease inhibitor cocktail tablet (Roche). The lysate was clarified by centrifuging at 264,900 × g for 1 hr. The clarified supernatant was incubated with IgG Sepharose beads (GE Healthcare Life Sciences) for 1 hr. The beads were transferred to a gravity flow column, washed with Dynein-lysis buffer supplemented with 250 mM potassium chloride, 0.1 mM Mg-ATP, 0.5 mM Pefabloc and 0.1% Triton, and with TEV buffer (10 mM Tris–HCl [pH 8.0], 150 mM potassium chloride, 10% glycerol, 1 mM DTT, and 0.1 mM Mg-ATP). Dynein was cleaved from IgG beads via incubation with 0.15 mg/mL TEV protease (purified in the Reck-Peterson lab) overnight at 4°C. Cleaved dynein was concentrated using 100K MWCO concentrator (EMD Millipore), filtered by centrifuging with Ultrafree-MC VV filter (EMD Millipore) in a tabletop centrifuge and flash frozen in liquid nitrogen.

##### S. cerevisiae Lis1

*S. cerevisiae* Lis1 was purified from *S. cerevisiae* using 8xHis and ZZ tags as previously described^41^. In brief, liquid-nitrogen frozen pellets were ground in a pre-chilled coffee grinder, resuspended in buffer A (50 mM potassium phosphate [pH 8.0], 150 mM potassium acetate, 150 mM sodium chloride, 2 mM magnesium acetate, 5 mM β-mercaptoethanol, 10% glycerol, 0.2% Triton, 0.5 mM Pefabloc) supplemented with 10 mM imidazole (pH 8.0) and cOmplete EDTA-free protease inhibitor cocktail tablet, and spun at 118,300 x g for 1h. The clarified supernatant was incubated with Ni-NTA agarose (QIAGEN) for 1 hr. The Ni beads were transferred to a gravity column, washed with buffer A + 20 mM imidazole (pH 8.0), and eluted with buffer A + 250 mM imidazole (pH 8.0). The eluted protein was incubated with IgG Sepharose beads for 1 hr. IgG beads were transferred to a gravity flow column, washed with buffer A + 20 mM imidazole (pH 8.0) and with modified TEV buffer (50 mM Tris–HCl [pH 8.0], 150 mM potassium acetate, 2 mM magnesium acetate, 1 mM EGTA, 10% glycerol, 1 mM DTT). Lis1 was cleaved from the IgG beads by the addition of 0.15 mg/mL TEV protease (purified in the Reck-Peterson lab) for 1 hr at 16°C. Cleaved proteins were filtered by centrifuging with Ultrafree-MC VV filter (EMD Millipore) in a tabletop centrifuge and flash frozen in liquid nitrogen.

##### Human Dynein

Full-length human SNAP-dynein was expressed in Sf9 cells as described previously^24,27,69^. Briefly, frozen Sf9 cell pellets from 2x 600 mL culture were resuspended in 80 mL of Dynein-lysis buffer with 0.1 mM Mg-ATP, 0.5 mM Pefabloc, 0.05% Triton, and cOmplete EDTA-free protease inhibitor cocktail tablet (Roche) and lysed using a Dounce homogenizer (10 strokes with a loose plunger and 15 strokes with a tight plunger). The lysate was clarified by centrifuging at 183,960 x g for 88 min in a Type 70 Ti rotor (Beckman). The clarified supernatant was incubated with 4 mL of IgG Sepharose 6 Fast Flow beads (GE Healthcare Life Sciences) for 3-4 hours on a roller. The beads were transferred to a gravity flow column, washed with 200 mL of Dynein-lysis buffer and 300 mL of TEV buffer (50 mM Tris– HCl [pH 8.0], 250 mM potassium acetate, 2 mM magnesium acetate, 1 mM EGTA, 1 mM DTT, 0.1 mM Mg-ATP, 10% (v/v) glycerol). For fluorescent labeling of carboxy-terminal SNAPf tag, dynein-coated beads were labeled with 5 µM SNAP-Cell-TMR (New England Biolabs) in the column for 10 min at room temperature and unbound dye was removed with a 300 mL wash with TEV buffer at 4°C. The beads were then resuspended and incubated in 15 mL of TEV buffer supplemented with 0.5 mM Pefabloc and 0.2 mg/mL TEV protease (purified in the Reck-Peterson lab) overnight on a roller. The supernatant containing cleaved protein was concentrated using a 100K MWCO concentrator (EMD Millipore) to 500 µL and purified via size exclusion chromatography on a TSKgel G4000SWXL column (TOSOH Bioscience) with GF150 buffer (25 mM HEPES [pH7.4], 150 mM potassium chloride, 1mM magnesium chloride, 5 mM DTT, 0.1 mM Mg-ATP) at 1 mL/min. The peak fractions were collected, buffer exchanged into a GF150 buffer supplemented with 10% glycerol, concentrated to 0.1-0.5 mg/mL using a 100K MWCO concentrator (EMD Millipore). Purity was evaluated on SDSPAGE gels and protein aliquots were snap frozen in liquid N2 and stored at -80°C.

##### Human dynactin

Dynactin (p62-HaloTag-3XFLAG) was purified from a stable 293T cell line as previously described^24,26,27^. Briefly, frozen pellets from 293T cells (160 × 15 cm plates) were resuspended in Dynein-lysis buffer supplemented with 0.1 mM Mg-ATP, 0.5 mM Pefabloc, 0.05% Triton, and cOmplete EDTAfree protease inhibitor cocktail tablet (Roche). and gently mixed at 4°C for 15 min. The lysed cells were then centrifuged at 66,000 x rpm in a Ti70 rotor (Beckman) at 4°C for 30 min. The clarified lysate was retrieved and added to 3 mL packed anti-FLAG M2 agarose resin (Sigma) and incubated with gentle mixing at 4°C for 16 hours. After incubation, the lysate/resin mixture was centrifuged at 1000 x g for 2 minutes at 4°C to pellet the resin and the supernatant was decanted. The resin was transferred to a column at 4°C and the column was washed with 100 mL low salt wash buffer (30 mM HEPES, pH 7.4; 50 mM KOAc; 2 mM MgOAc; 1 mM EGTA, pH 7.5; 10% glycerol; 1 mM DTT; 0.5 mM ATP; 0.5 mM Pefabloc; 0.02% Triton X-100), 100 mL high salt wash buffer (30 mM HEPES, pH 7.4; 250 mM KOAc; 2 Mm MgOAc; 1 mM EGTA, pH 7.5; 10% glycerol; 1 mM DTT; 0.5 mM ATP; 0.5 mM Pefabloc; 0.02% Triton X-100), and finally with 50 mL of low salt wash buffer. The resin was resuspended in 800 µL of low salt wash buffer containing 2 mg/mL 3X-FLAG peptide (ApexBio) and incubated for 30 min at 4°C. The mixture was retrieved and centrifuged through a small filter column to remove the resin. The eluate was then loaded onto a Mono Q 5/50 GL 1 mL column on an AKTA FPLC (GE Healthcare). The column was washed with 5 mL of Buffer A (50 mM Tris-HCl, pH 8.0; 2 mM MgOAc; 1 mM EGTA; 1 mM DTT) and then subjected to a 26 mL linear gradient from 35-100% Buffer B mixed with Buffer A (Buffer B = 50 mM Tris-HCl, pH 8.0; 1 M KOAc; 2 mM MgOAc; 1 mM EGTA; 1 mM DTT), followed by 5 mL additional 100% Buffer B. Fractions containing pure dynactin (∼75-80% Buffer B) were pooled and buffer exchanged through iterative rounds of dilution and concentration on a 100 kDa MWCO centrifugal filter (Amicon Ultra, Millipore) using GF150 buffer with 10% glycerol. Purity was evaluated on SDS-PAGE gels and protein aliquots were snap frozen in liquid N2 and stored at -80°C.

##### Human Lis1

Lis1 constructs were purified from frozen sf9 cell pellets from a 600 mL culture as described previously^69^. Lysis and clarification steps were similar to full-length dynein purification except lysis was performed in Lis1-lysis buffer (30 mM HEPES [pH 7.4], 50 mM potassium acetate, 2 mM magnesium acetate, 1 mM EGTA, 300 mM potassium chloride, 1 mM DTT, 0.5 mM Pefabloc, 10% (v/v) glycerol) supplemented with cOmplete EDTA-free protease inhibitor cocktail tablet (Roche) per 50 mL was used. The clarified supernatant was incubated with 0.5 mL of IgG Sepharose 6 Fast Flow beads (GE Healthcare Life Sciences) for 2-3 hours on a roller. The beads were transferred to a gravity flow column, washed with 20 mL of Lis1-lysis buffer, 100 mL of modified TEV buffer (10 mM Tris–HCl [pH 8.0], 2 mM magnesium acetate, 150 mM potassium acetate, 1 mM EGTA, 1 mM DTT, 10% (v/v) glycerol) supplemented with 100 mM potassium acetate, and 50 mL of modified TEV buffer. Lis1 was cleaved from IgG beads via incubation with 0.2 mg/mL TEV protease overnight on a roller. The cleaved Lis1 was filtered by centrifuging with an Ultrafree-MC VV filter (EMD Millipore) in a tabletop centrifuge. Purity was evaluated on SDS-PAGE gels and protein aliquots were snap frozen in liquid N_2_ and stored at -80°C.

##### Human BICD2

BICD2 construct containing amino-terminal sfGFP was expressed and purified as previously described^26^. In brief, BL-21[DE3] cells (New England iolabs) were grown at optical density at 600 nm of 0.4–0.6 and protein expression was induced with 0.1 mM IPTG for 16h at 18 °C. Frozen cell pellet from a 2 L culture was resuspended in 60 ml of lysis buffer (30 mM HEPES pH7.4, 50 mM potassium acetate, 2mM magnesium acetate, 1mM EGTA, 1mM DTT and 0.5mM Pefabloc, 10% (v/v) glycerol) supplemented with cOmplete EDTA-free protease inhibitor cocktail tablet (Roche) per 50 ml and 1 mg/ml lysozyme. The resuspension was incubated on ice for 30 min and lysed by sonication. The lysate was clarified by centrifuging at 66,000g for 30min in Type 70 Ti rotor (Beckman). The clarified supernatant was incubated with 2ml of IgG Sepharose 6 Fast Flow beads (GE Healthcare Life Sciences) for 2 h on a roller. The beads were transferred into a gravity-flow column, washed with 100ml of activating-adaptor-lysis buffer supplemented with 150 mM potassium acetate and 50 ml of cleavage buffer (50 mM Tris-HCl pH 8.0, 150 mM potassium acetate, 2 mM magnesium acetate, 1 mM EGTA, 1 mM DTT, 0.5 mM Pefabloc and 10% (v/v) glycerol). The beads were then resuspended and incubated in 15 ml of cleavage buffer supplemented with 0.2 mg/ml TEV protease overnight on a roller. The supernatant containing cleaved protein was concentrated using a 50 kDa MWCO concentrator (EMD Millipore) to 1 ml, filtered by centrifuging with Ultrafree-MC VV filter (EMD Millipore) in a tabletop centrifuge, diluted to 2 ml in buffer A (30 mM HEPES pH 7.4, 50 mM potassium acetate, 2 mM magnesium acetate, 1 mM EGTA, 10% (v/v) glycerol and 1mM DTT) and injected into a MonoQ 5/50 GL column (GE Healthcare and Life Sciences) at 1 ml/min. The column was prewashed with 10 CV of buffer A, 10 CV of buffer B (30 mM HEPES pH7.4, 1 M potassium acetate, 2 mM magnesium acetate, 1 mM EGTA, 10% (v/v) glycerol and 1 mM DTT) and again with 10 CV of buffer A at 1 ml/min. To elute, a linear gradient was run over 26 CV from 0–100% buffer B. The peak fractions containing sfGFP-BICD2s were collected and concentrated using a 50 kDa MWCO concentrator (EMD Millipore) to 0.2 ml. Protein was then diluted to 0.5 ml in GF150 buffer and further purified using size-exclusion chromatography on a Superose 6 Increase 10/300GL column (GE Healthcare and Life Sciences) with GF150 buffer at 0.5 ml/min. The peak fractions were collected, buffer-exchanged into a GF150 buffer supplemented with 10% glycerol, concentrated to 0.2–1 mg/ml using a 50 kDa MWCO concentrator (EMD Millipore) and flash-frozen in liquid nitrogen

#### TIRF microscopy

Imaging was performed with an inverted microscope (Nikon, Ti-E Eclipse) equipped with a 100x 1.49 N.A. oil immersion objective (Nikon, Plano Apo). The xy position of the stage was controlled by ProScan linear motor stage controller (Prior). The microscope was equipped with an MLC400B laser launch (Agilent) equipped with 405 nm (30 mW), 488 nm (90 mW), 561 nm (90 mW), and 640 nm (170 mW) laser lines. The excitation and emission paths were filtered using appropriate single bandpass filter cubes (Chroma). The emitted signals were detected with an electron multiplying CCD camera (Andor Technology, iXon Ultra 888). Illumination and image acquisition was controlled by NIS Elements Advanced Research software (Nikon).

#### Single-molecule motility assays

Single-molecule motility assays were performed in flow chambers using the TIRF microscopy set up described above. To reduce non-specific binding biotinylated and PEGylated coverslips (Microsurfaces) with microtubules polymerized from tubulin prepared from bovine brain as previously described18. Microtubules contained ∼10% biotin-tubulin to allow for attachment to streptavidin-coated cover slip and ∼10% Alexa Fluor 488 (Thermo Fisher Scientific) tubulin for visualization. Imaging was done in Dynein-lysis buffer supplemented with 20 µM taxol, 1 mg/mL casein, 5 mM Mg-ATP, 71.5 mM βME (beta mercaptoethanol) and an oxygen scavenger system containing, 0.4% glucose, 45 μg/ml glucose catalase (Sigma-Aldrich), and 1.15 mg/ml glucose oxidase (Sigma-Aldrich). Images were recorded every 0.3 sec for 3 minutes. Movies showing significant drift were not analyzed.

All movies were collected by measuring TMR-dynein signal with the following protein concentrations: 83.5 pM TMR-dynein, 665 pM unlabeled dynactin, 5 nM BICD2, and 300 nM Lis1. For conditions missing Lis1, the corresponding matching buffer was used. The dynein, dynactin, and BICD2 complexes were incubated on ice for 10 minutes prior to Lis1 or buffer addition. Each protein mixture was then incubated on ice for an additional 10 minutes prior to TIRF imaging.

#### TIRF data analysis

The velocity of moving particles was calculated from kymographs generated in Fiji as described previously^52^. Velocities were only calculated from molecules that moved processively for greater than 5 frames. Non-motile or diffusive events were not considered in velocity calculations. Processive events were defined as events that move unidirectionally and do not exhibit directional changes greater than 600 nm. Diffusive events were defined as events that exhibit at least one bidirectional movement greater than 600 nm in each direction. Single-molecule movements that change apparent behavior (e.g., shift from non-motile to processive) were considered as multi-velocity events and counted as multiple events. Landing rates were calculated by counting the number of processive events for each microtubule in individual movies and dividing this number by the microtubule length. Data visualization and statistical analyses were performed using GraphPad Prism (9.2; GraphPad Software), Excel (version 16.52; Microsoft), and Fiji (2.0)^67^. Brightness and contrast were adjusted in Fiji for all videos and kymographs. The exact value of n, evaluation of statistical significance, and the specific statistical analysis are described in the corresponding figures and figure legends. All experiments were analyzed from at least three independent replicates.

**Supplementary Video 1. The structure of *S. cerevisiae* “Chi” (dynein-Lis1)**.

The video shows the structure of Chi, highlighting the different interfaces involving Lis1: those between Lis1 and dynein at site_ring_ and site_stalk_, the one between Lis1 at site_ring_ and the opposite dynein monomer responsible for the formation of Chi, and the one that mediates the Lis1-Lis1 interaction. The residues on Lis1 that were mutated in this study are highlighted.

**Extended Figure 1.**
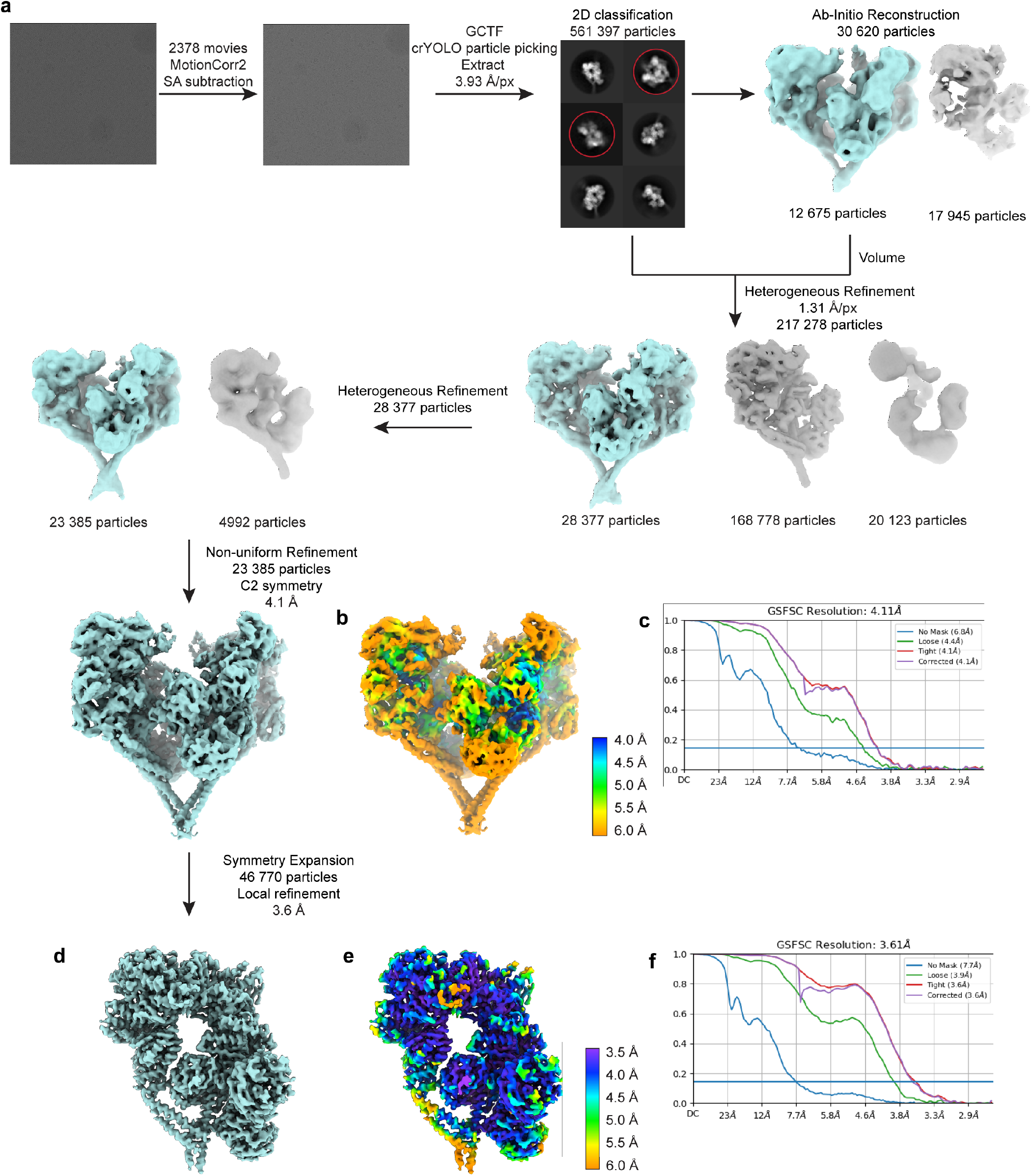
Cryo-EM data processing workflow. **a**. Processing workflow for Chi. Particles belonging to teal maps were carried into the next round of processing. **b**. Cryo-EM map of Chi (dyneinE2488Q-Lis1)2 colored by local resolution. **c**. Fourier shell correlation (FSC) plot for Chi. **d**. Symmetry expansion followed by local refinement map of dyneinE2488Q-(Lis1)2. **e**. Local refinement of symmetry expanded dyneinE2488Q-(Lis1)2 colored by local resolution. **f**. FSC plot for local refinement map.

**Extended Figure 2.**
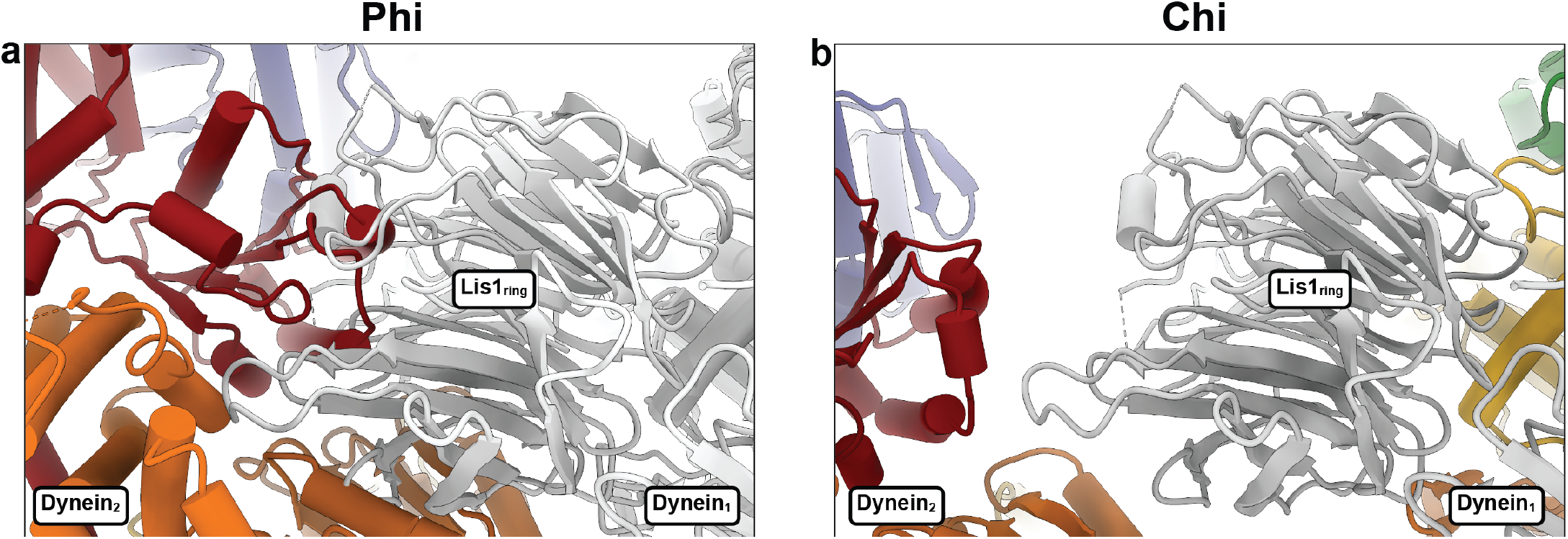
Binding of Lis1 to dynein is sterically incompatible with Phi. **a**. Phi dynein (PDB 5NUG) was aligned to dynein1 in Chi, which is shown in grey in this panel. Phi’s dynein_2_, shown in color, clashes with the Lis1 bound at site_ring_ in Chi’s dynein_1_. **b**. Close up of our model of Chi equivalent to the view in (a), showing Lis1 bound at site_ring_ on dynein_1_. Part of dynein_2_, with which the Lis1 at site_ring_ interacts in Chi, is also shown.

**Extended Figure 3.**
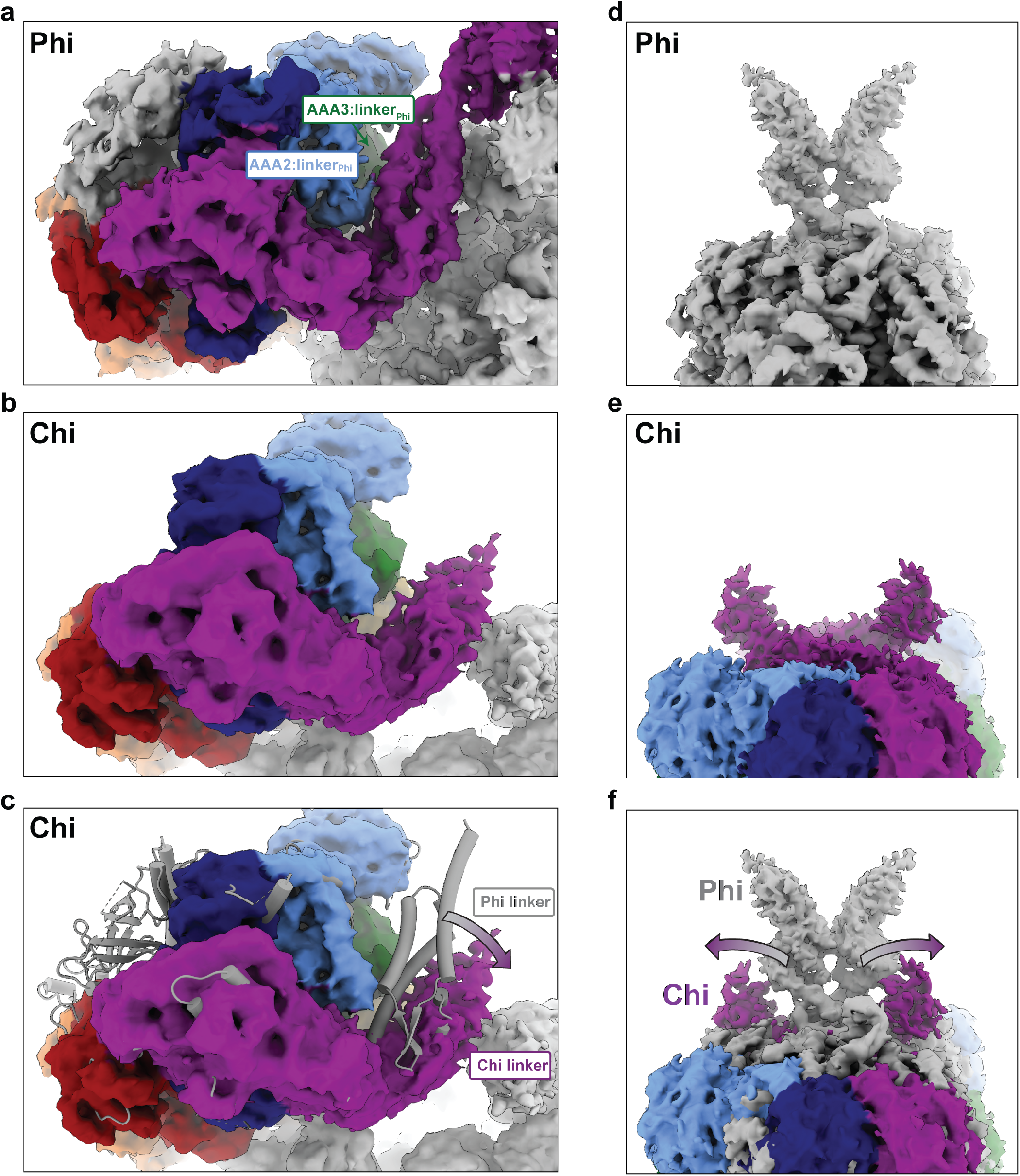
The conformation of the linker is different in Chi and Phi. **a**. The linker docks onto AAA2 and AAA3 in Phi. b. In contrast, the linker is disengaged from the AAA ring in Chi. **c**. Overlay of Phi (a) and Chi (b) highlights the different positions of the linker. **d, e**. The N-termini of the linker adopt different paths in Phi (d) and Chi (e). **f**. Overlay of Phi (d) and Chi (e) shows how the linkers are pulled apart in Chi.

**Extended Figure 4.**
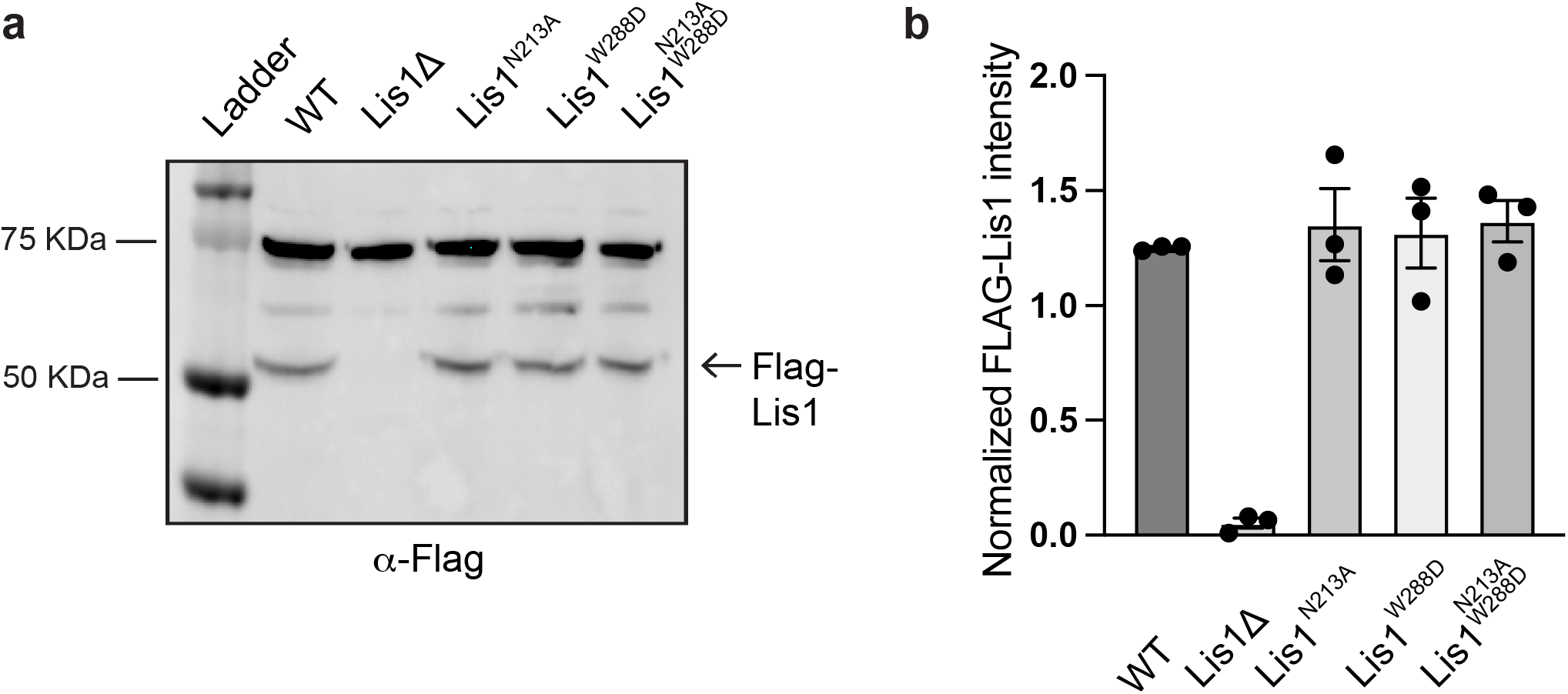
Western blot analysis of Lis1 mutant expression. **a**. Cell lysates from wild type (WT), Lis1Δ or mutant Lis1 (Lis1^N213A^, Lis1^W288D^ and Lis1^N213A, W88D^) *S. cerevisiae* strains containing Flag-tagged Lis1 (Pac1) were immunoprecipitated using anti-Flag agarose beads and immunoblotted for the FLAG peptide (α-FLAG). **b**. Quantitative analysis of Lis1 expression (α-FLAG band) normalized against the non-specific band observed at ∼75 KDa for three independent immunoprecipitations.

**Extended Figure 5.**
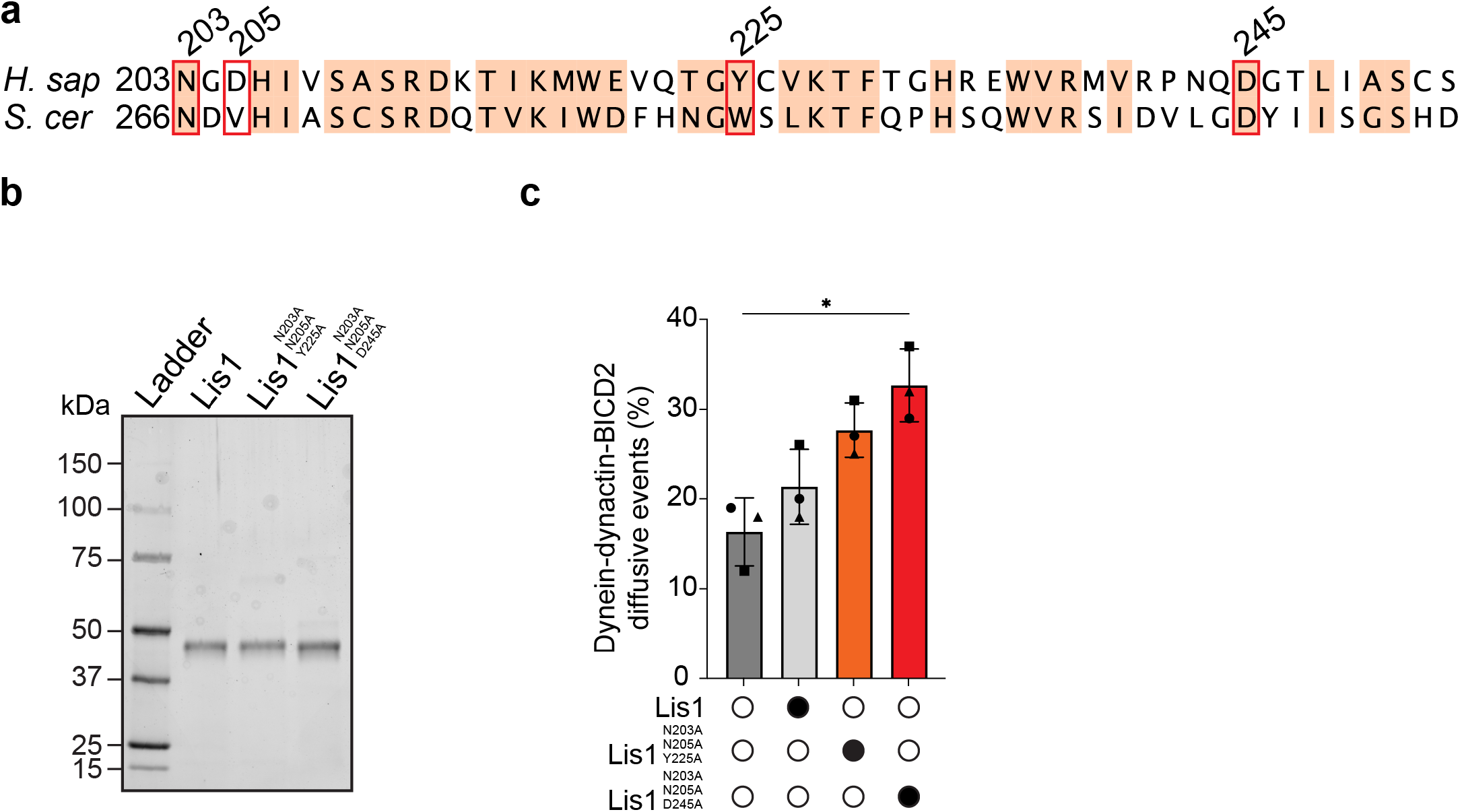
Chi mutations in human Lis1. **a**. Sequence conservation between *S. cerevisiae* (*S. cer*) and human (*H. sap*) Lis1 in the vicinity of the dynein binding site in the Chi structure. Residues with 70% conservation or higher are shaded in light red. The residues mutated in this study: N203, N205, Y225, and D245 are highlighted by red boxes. **b**. Sodium dodecyl sulfate–polyacrylamide gel electrophoresis (SDS–PAGE) of the human Lis1 constructs used for motility assays. **c**. Comparison of percent diffusive events ± SEM from single-molecule TMR-dynein-dynactin-BICD2 assays performed in the absence (white circle) or presence (black circle) of different Lis1 constructs. The data points are represented as triangles, circles, and squares, corresponding to single measurements within each technical replicate (no Lis1, n = 52; Lis1, n = 29; Lis1^N203A, D205A, Y225A^, n = 38; Lis1^N203A, D205A, D245A^, n = 43). ns, not significant, * < 0.05. One-Way Anova with Brown-Forsythe test.

**Supplementary Table 1.** S. cerevisiae strains used in this study. *DHA* and *SNAP* refer to the HaloTag (Promega) and SNAP-tag (NEB), respectively. TEV indicates a Tev protease cleavage site. P_GAL1_ denotes the galactose promoter, which was used for inducing strong expression of Lis1 and dynein motor domain constructs. Amino acid spacers are indicated by g (glycine) and gs (glycine-serine).

| Strain | Genotype | Source |
| --- | --- | --- |
| RPY1 | W303a (*MATa; his3-11,15; ura3-1; leu2-3,112; ade2-1; trp1-1*) | (Eshel et al., 1993) |
| RPY816 | W303a; *pep4Δ::HIS5; prb1Δ; GAL1-8HIS-ZZ-Tev-PAC1; dyn1Δ::CgLEU2; ndl1Δ::HPH* | (Huang et al., 2012) |
| RPY1654 | W303a; *pep4Δ::HIS5; prb1Δ; PAC11-13xMYC-TRP1; PGAL1-ZZTev-DYN1(331kDa) E2488Q; pac1Δ::HygroR* | (DeSantis et al., 2017) |
| RPY1385 | *MATa lys2-801 leu2-Δ1 his3-Δ200 trp1-Δ63 DYN1-3XGFP::TRP1; ura3-52::CFP-TUB1::URA3; SPC110-tdTomato::SpHIS5; ura3Δ::KanMX* | (DeSantis et al., 2017) |
| RPY1717 | *MATa lys2-801; leu2-Δ1; his3-Δ200; trp1-Δ63; DYN1-3XGFP::TRP1; ura3-52::CFP-TUB1::URA3; SPC110-tdTomato::SpHIS5; ura3Δ::KanMX; pac1Δ::KIURA3* | (DeSantis et al., 2017) |
| RPY1795 | *MATa lys2-801; leu2-Δ1; his3-Δ200; trp1-Δ63; DYN1-3XGFP::TRP1; ura3-52::CFP-TUB1::URA3; SPC110-tdTomato::SpHIS5; ura3Δ::KanMX; Flag-PAC1* | (Gillies et al., 2022) |
| RPY1814 | *MATa lys2-801; leu2-Δ1; his3-Δ200; trp1-Δ63; DYN1-3XGFP::TRP1; ura3-52::CFP-TUB1::URA3; SPC110-tdTomato::SpHIS5; ura3Δ::KanMX; Flag-pac1(W288D)* | This work |
| RPY1831 | *MATa lys2-801; leu2-Δ1; his3-Δ200; trp1-Δ63; DYN1-3XGFP::TRP1; ura3-52::CFP-TUB1::URA3; SPC110-tdTomato::SpHIS5; ura3Δ::KanMX; Flag-pac1(N213A)* | This work |
| RPY1834 | *MATa lys2-801; leu2-Δ1; his3-Δ200; trp1-Δ63; DYN1-3XGFP::TRP1; ura3-52::CFP-TUB1::URA3; SPC110-tdTomato::SpHIS5; ura3Δ::KanMX; Flag-pac1(N213A-W288D)* | This work |

## Notes

### Competing Interest Statement

The authors have declared no competing interest.

